# Castration alters the circadian clock and *Ucp1* expression in subcutaneous white adipose tissue of mice

**DOI:** 10.1101/2022.06.13.495888

**Authors:** Tsuyoshi Otsuka, Hiroki Onishi, Mao Suzuki, Mami Marusawa

**Author notes:** **Corresponding author’s email address** Tsuyoshi Otsuka.

## Abstract

Castration increases the risk of metabolic syndromes in mammals. In addition, recent studies have demonstrated that castration increases the expression of *Ucp1* in subcutaneous white adipose tissue (scWAT). However, how castration leads to thermogenic aberration remains poorly understood. Here, we found that castrated mice showed elevated *Ucp1* expression in scWAT in response to cold exposure with an increase in body temperature in the light phase and a reduction in body weight gain with fat reduction despite showing no alterations in food intake, locomotor activity in a novel environment, and gastrocnemius muscle weight. We also analyzed the circadian behavioral rhythm and the expression rhythm of clock genes in the suprachiasmatic nucleus (SCN) and scWAT. We found that castrated mice showed fluctuations in circadian locomotor activity and a decrease in the expression rhythm of clock genes in the SCN (*clock*) and scWAT (*Bmal1* and *Per2*). Moreover, the expression of *Ucp1* in scWAT was rhythmic, and it was increased in castrated mice, while the expression of clock genes was decreased. These results indicate that castration may impact browning in scWAT through variations in the circadian clock and *Ucp1* expression.

## Introduction

Castration is generally performed on livestock and companion animals to control aggressive behaviors, improve meat quality, prevent unwanted breeding, and prevent estrus. Castration in animals provides benefits to both humans and coexisting animals but has the disadvantages of promoting the development of diabetes, adiposity, and hypothyroidism in animals [1–3]. In humans, although sex reassignment surgery is performed to solve the problem of gender dysphoria, it is well known that there are several drawbacks, such as obesity, insulin resistance and dyslipidemia [4–6]. Furthermore, castrated mice fed a high-fat diet exhibit abdominal obesity with intestinal microbiome alterations [7,8]. However, interestingly, it has been reported that castrated mice lose weight under a normal diet compared to control mice [7,9]. Additionally, castrated and androgen receptor-knockout mice show reduced epididymal fat and subcutaneous adipocyte size and an associated reduction in body fat percentage [10]. These reports in several animals and men suggest that although the pathogenic mechanism of castration has not yet been elucidated, castration affects the lipid metabolic capacity of all animals and triggers a variety of metabolic disorders.

Circadian rhythms are controlled by the transcriptional-translational feedback loop system of several clock genes. Heterodimers of the CLOCK and BMAL1 proteins bind to the E-box and promote the expression of *Per* and *Cry*, which act as their own repressors. CLOCK and BMAL1 proteins also promote *Rev-erbα* and *Ror*, which are involved in the regulation of *Bmal1* transcription [11,12]. In previous studies, it has been demonstrated that these circadian clock systems regulate various lipid metabolic functions and that their disturbance leads to metabolic disorders [13,14]. In rodents, clock gene mutant mice exhibit obesity, metabolic syndrome, dyslipidemia, and ectopic fat formation [15,16]. A high-fat diet in healthy mice leads to an increase in body weight with the disturbance of circadian rhythms [17]. Interestingly, various studies have demonstrated that castration disrupts the circadian clock in addition to altering lipid metabolic capacity in mice, as mentioned above. In male and female mice, circadian behavioral rhythms disappear after gonadectomy [18]. Moreover, castration causes diminished *Bmal1* and *Per2* circadian expression rhythm in prostate tissue [19]. Therefore, we hypothesized based on these associations that castration-induced alterations in lipid metabolism are related to circadian clock fluctuations. In this study, we focused on heat production in adipocytes as a target of changes in lipid metabolism. Recently, it has become clear that beige adipocytes are deeply associated with lipid metabolism and energy expenditure, in addition to brown adipose tissue (BAT). Beige adipocytes are known to differentiate from white adipose tissue (WAT) that is stimulated by cold exposure or administration of β3-adrenergic receptor agonists, and these stimuli lead to the expression of genes, such as *Pgc1-α, Pparγ*, and *Prdm16*, in WAT that then induce the expression of the thermogenic gene *Ucp1* [20–22]. Therefore, we investigated how castration affects the circadian clock and heat production in WAT and BAT of mice.

## Materials and methods

### Animals

In all experiments, C57BL/6J male mice were obtained from Japan SLC (Shizuoka, Japan). These animals were housed in a light-dark control box (150 W × 45 D × 48 H) so that they were not affected by light from the external environment. All mice in the box were maintained under a 12-h light/12-h dark (LD) cycle at a temperature of 23 ± 2 °C. A standard diet for laboratory rodents (MF, Oriental Yeast, Tokyo, Japan) and fresh water were available *ad libitum*. Castration was performed under anesthesia (2 mg ketamine + 0.2 mg xylazine with saline/1 mouse), a small incision of approximately 1.5 cm was made in the midline of the lower abdomen, and vas deferens ligation was performed in approximately 6–8-week-old mice. For sham surgery, only a lower abdominal incision was performed under anesthesia. After an approximately 1–2-week acclimation period after surgery, sham-operated (Sham) mice and castrated (Cast) mice were housed individually, and body weight and food intake were calculated for an 8-week period. After the experimental period, the mice were sacrificed by isoflurane overanesthesia, and the gastrocnemius muscle weight and epididymal fat (eFAT) weight were measured. All experimental mice were housed and experiments were performed in accordance with the rules stipulated by the Gifu University animal experiment handling regulations.

### Behavioral tests

To measure locomotor activity in a novel environment, after an approximately 1–2-week acclimation period after surgery, we performed an open field test. Mice were placed in the center of the field (40 W × 40 D × 40 H) and allowed to move freely for 5 min. The behavior was recorded and analyzed by dividing the field into 25 squares (5 × 5 grid lines). The number of grid lines crossed was counted on the PC monitor.

To assess the circadian behavioral rhythm, male mice after 1∼2 weeks of acclimation post-surgery were housed individually in home cages, and circadian locomotor activity was monitored (AS – 10F, MELQUEST, Toyama, Japan) under a 10-day LD cycle and 20-day constant darkness (DD) cycle. This monitoring system detects the radiated body heat and quantifies the animals’ movement. Data were continuously recorded in 2-minute bins with a date collection (CIF4, Actmaster4, MELQUEST, Toyama, Japan).

### Core body temperature rhythm measurements

The core body temperature was measured using a temperature data logger (Thermo manager data logger, KN laboratories, Osaka, Japan). At the time of castration or sham surgery, a data logger was implanted in 6–8-week-old mice, 1∼2 weeks of acclimation time was provided, and then, 2 days of measurement was performed.

### Tissue collection

To obtain tissue samples, after 1∼2 weeks of recovery from surgery, all mice were sacrificed by decapitation under isoflurane anesthesia. Samples for confirming the expression of heat production-related genes were obtained at Zeitgeber time (ZT) 6 after cold stimulation at 4 °C for 4 hours (ZT 2∼6). Twenty-four-hour rhythm samples were sampled at ZT 2, 6, 10, 14, 18, and 22 (ZT 0 = light on and ZT 12 = light off). Decapitation under dark conditions was performed under dim red light. BAT, scWAT, and the suprachiasmatic nucleus (SCN) were sampled from dissected mice. In SCN sampling, brains were cut using a cooled mouse brain matrix (Brainscience idea Co., Ltd., Osaka, Japan), and the SCN was removed carefully from its cut surface using extrafine tweezers.

### RNA extraction and quantitative RT–PCR

To analyze various gene expression levels, total RNA from samples was extracted with TRIzol Isolation Reagent (Thermo Fisher Scientific, Inc., Waltham, MA, USA). First-strand cDNA was synthesized from 250 ng of RNA using a ReverTra Ace qPCR RT kit (TOYOBO, Osaka, Japan). Quantitative RT–PCR was performed using the Mx3000P Real-Time QPCR System (Agilent Technologies, Tokyo, Japan). Brilliant III Ultra-Fast SYBR Green QPCR Master Mix (Agilent Technologies, Tokyo, Japan), 250 nM of each primer, and cDNA in a 10-μl volume were dispensed into the well. The PCR conditions were 95 °C for 180 seconds, followed by 35 cycles of 95 °C for 5 seconds and 60 °C for 20 seconds. Gene expression levels were calculated by the comparative CT method using *Gapdh* as the housekeeping gene.

### Statistical analysis

For all the data, the results are indicated as the mean ± SEM. Student’s *t* test was used to assess the significant difference between two groups. For comparison of multiple groups, two-way ANOVA and subsequent Tukey–Kramer tests were applied to determine statistical significance. Values were considered significantly different at *p* < 0.05.

## Results

### Castration affects the expression of the thermogenic gene *Ucp1* and alters body weight

To investigate the effect of castration on the metabolic function of mice, we first measured food intake and body weight for 8 weeks. As a result, there was no significant difference in the amount of food intake per body weight between the Sham and Cast groups (Fig. 1A). However, the rate of weight gain decreased in the Cast group compared to the Sham group, as in previous reports [7,23] (Fig. 1B). When the visceral fat and muscle weights were measured to determine the cause of the weight loss in Cast mice, there was no significant difference in the gastrocnemius muscle weight, while the epididymal fat weight, which is the visceral fat, decreased in the Cast compared with the Sham group (Fig. C, D). Next, to examine the cause of the decrease in epididymal fat weight of Cast mice, locomotor activity was measured by the open field test. The result was that there was no significant difference in the number of lines crossed between the Sham and Cast mice (Fig. 1C). On the other hand, in the measurement of the core body temperature rhythm, Cast mice showed a higher body temperature than that of Sham mice (Fig. 1D). Notably, the Cast group had a high core body temperature in the light phase, which is the rest phase, and an increase in body temperature was observed without locomotor activity. For these reasons, it is believed that the decrease in the rate of weight gain was not due to an increase in the amount of spontaneous exercise but that the improvement in the heat production function accompanying the increase in body temperature had an effect. To confirm the reason for the increase in body temperature, the gene expression levels of the thermogenesis gene *Ucp1* and its related genes were measured in BAT and scWAT. In BAT, there was no significant difference in the expression of *Ucp1* and related genes between the Sham and Cast groups (Fig. 2A). In contrast, in scWAT, although there was no significant difference between Sham and Cast mice in the thermogenesis genes *Pgc1-α, Prdm16*, and *Pparγ*, the expression of the heat-producing gene *Ucp1* was significantly increased in Cast mice (Fig. 2B). These results indicate that castration increased the expression of *Ucp1* in the scWAT in mice and improved thermogenic function. Together, it is suggested that castration leads to the transition of scWAT into beige adipocytes, which produces a fat combustion effect by improving heat production and thereby reduces visceral fat.

**Figure 1.**
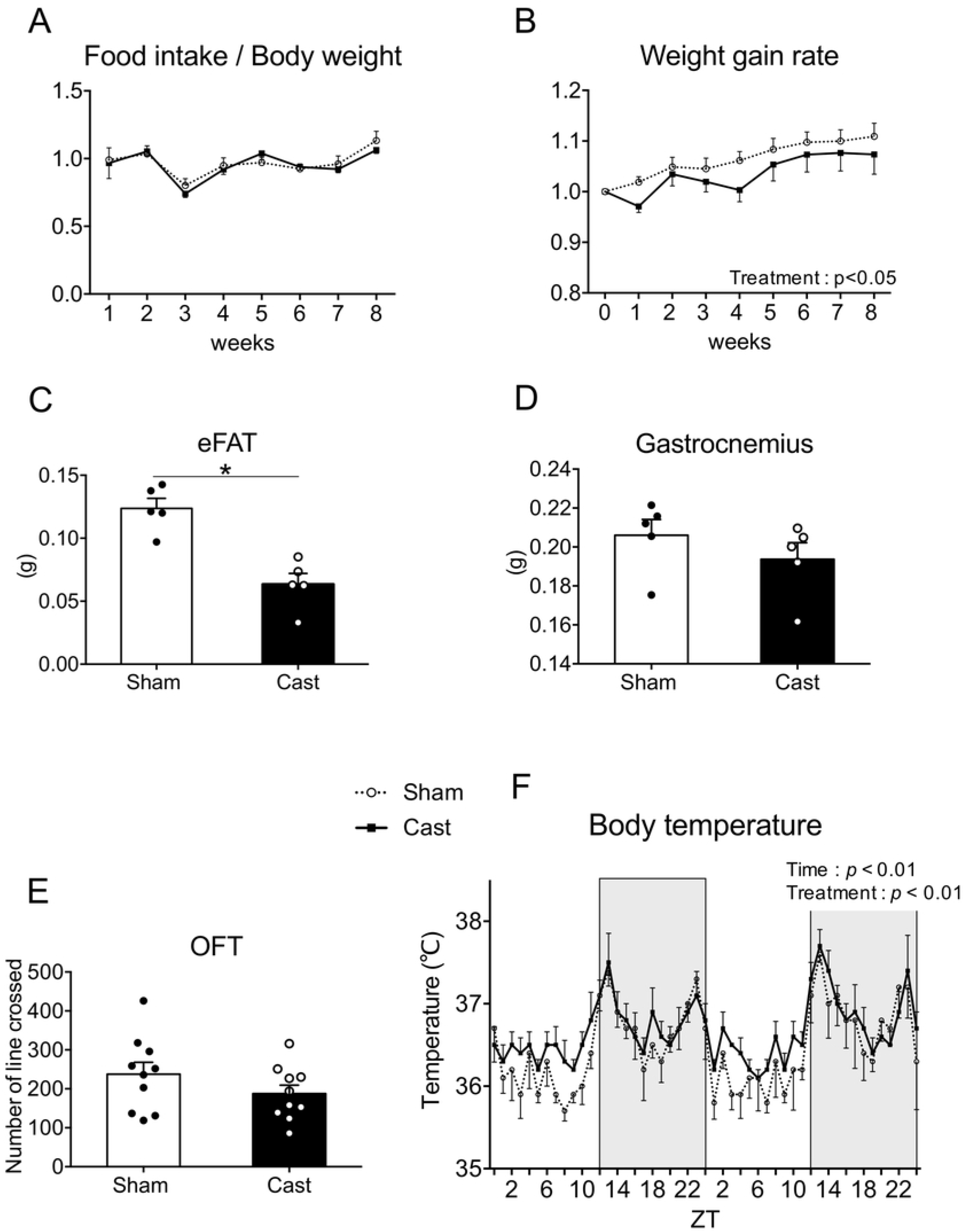
Effects of castration on food intake, body weight gain, body composition, behavior, and body temperature in mice. In approximately 6-to 8-week-old mice after 1∼2 weeks of acclimation, **(A)** the ratio of food intake to body weight and **(B)** body weight gain were measured for 8 consecutive weeks in Sham (dotted line) and Cast (solid line) mice (n = 5∼6). **(C)** eFAT and **(D)** gastrocnemius weight were measured after the Food intake/body weight ratio and body weight gain were measured (white bar: Sham, black bar: Cast) (n = 5). **(E)** Locomotor activity was measured by counting the line crossings in the OFT (n =10). **(F)** The circadian rhythm of body temperature was measured using a temperature data logger (n = 5). The bars at the bottom of the graph indicate the light period (white: ZT 0-12) and the dark period (black: ZT 12-24). The data are shown as the mean ± SEM. * *p* <0.05 (*t*-test). The main effects of time and treatment were detected by two-way ANOVA.

**Figure 2.**
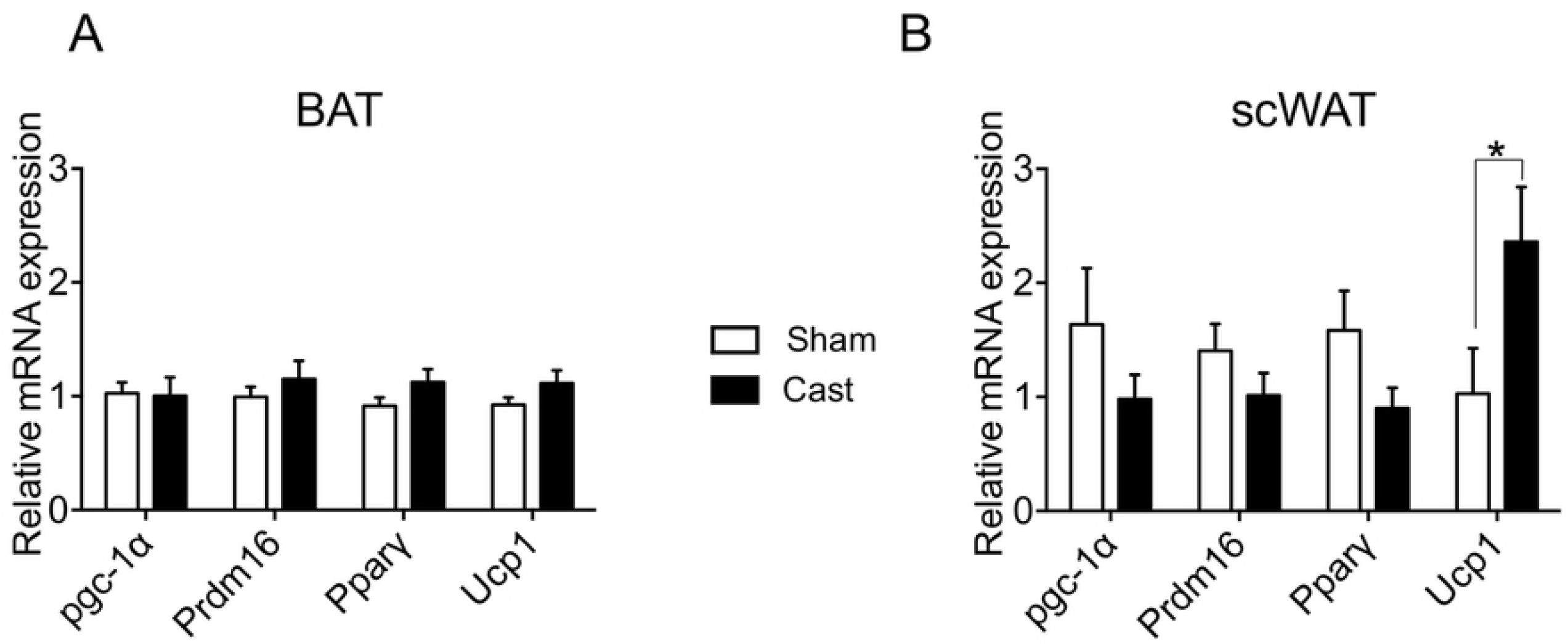
Expression of thermogenesis-related genes in BAT and scWAT. The relative expression of thermogenesis-related genes (*Pgc1-α, Prdm16, Ppar-γ, Ucp1*) in **(A)** BAT and **(B)** scWAT was analyzed at ZT 6 by quantitative RT-PCR (white bar: Sham, black bar: Cast) (n = 11∼12). The data are shown as the mean ± SEM. * *p* <0.05 (*t*-test).

### Castration disrupts the circadian clock in the SCN and scWAT and affects the *Ucp1* expression rhythm

To clarify the mechanism by which castration increases the expression of *Ucp1*, we focused on the circadian clock. Previous studies have shown that castration disrupts the circadian clock in prostate tissue [19]; however, it is unknown whether the central and scWAT circadian clocks are disrupted. To confirm the circadian clock rhythm after castration, we analyzed the daily locomotor activity. As a result of behavioral rhythm measurement, it was found that the castrated mice had decreased daily locomotor activity and disordered behavioral rhythm amplitude under DD conditions, although no significant differences were observed during the circadian period (Fig. 3A). Since it is well known that clock genes are deeply involved in the disturbance of the circadian clock, 24-h expression rhythms of clock genes in the SCN, which is the central region of the circadian clock, and the scWAT were measured. In the SCN, although other clock genes did not fluctuate significantly, the Cast group was found to have a significantly reduced *clock* expression level compared to that of the Sham group (Fig. 3B). In scWAT, the Cast group showed a significant decrease in *Bmal1* and *Per2* expression rhythm levels compared to those of the Sham group, and although there was no significant decrease in *Per1* expression, a decreasing tendency was found (Fig. 4A). Moreover, we measured the expression rhythm of *Ucp1* in scWAT. Consequently, it was found that the mouse scWAT had a rhythm, and Cast mice had a significantly increased *Ucp1* expression level compared to the level of Sham mice (Fig. 4B). These data imply that castration disorganizes the central and peripheral biological clocks and may also strongly affect the formation of *Ucp1* expression rhythm in scWAT.

**Figure 3.**
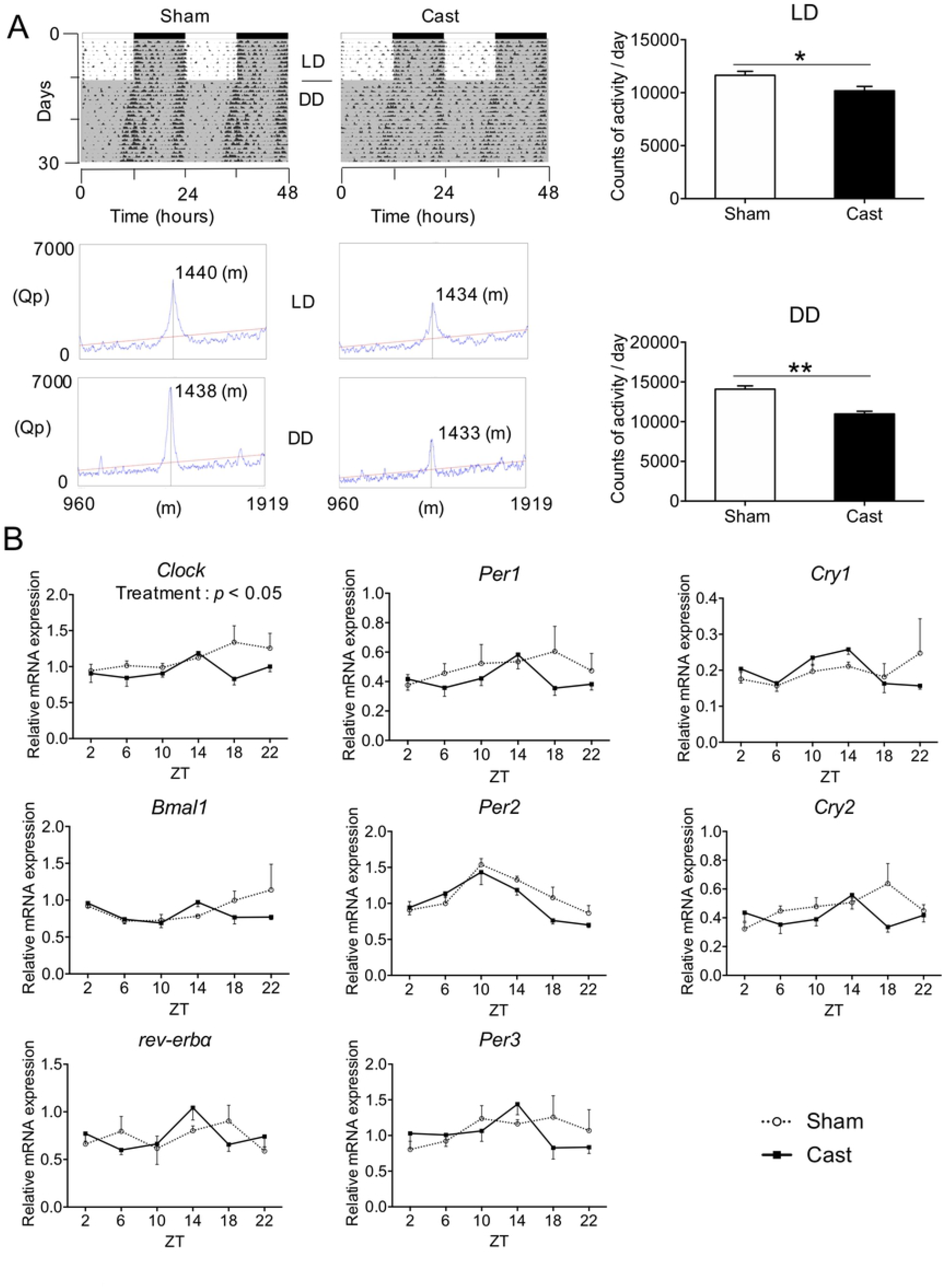
Effect of castration on the circadian activity and expression of clock genes in the SCN of mice. **(A)** Representative actograms, circadian periods, and amplitudes for Sham (left) and Cast (right) mice. The amount of daily activity was also analyzed under LD and DD cycles. **(B)** The relative expression of clock genes (*Clock, Bmal1, Rev-erbα, Per1, Per2, Per3, Cry1, Cry2*) was analyzed at ZT 2, 6, 10, 14, 18, and 22 by quantitative RT-PCR in the SCN of Sham (dot line) and Cast (solid line) mice (n = 4). The data are shown as the mean ± SEM. * *p* <0.05, ** *p* <0.01 (*t*-test). The main effect of treatment was detected by two-way ANOVA.

**Figure 4.**
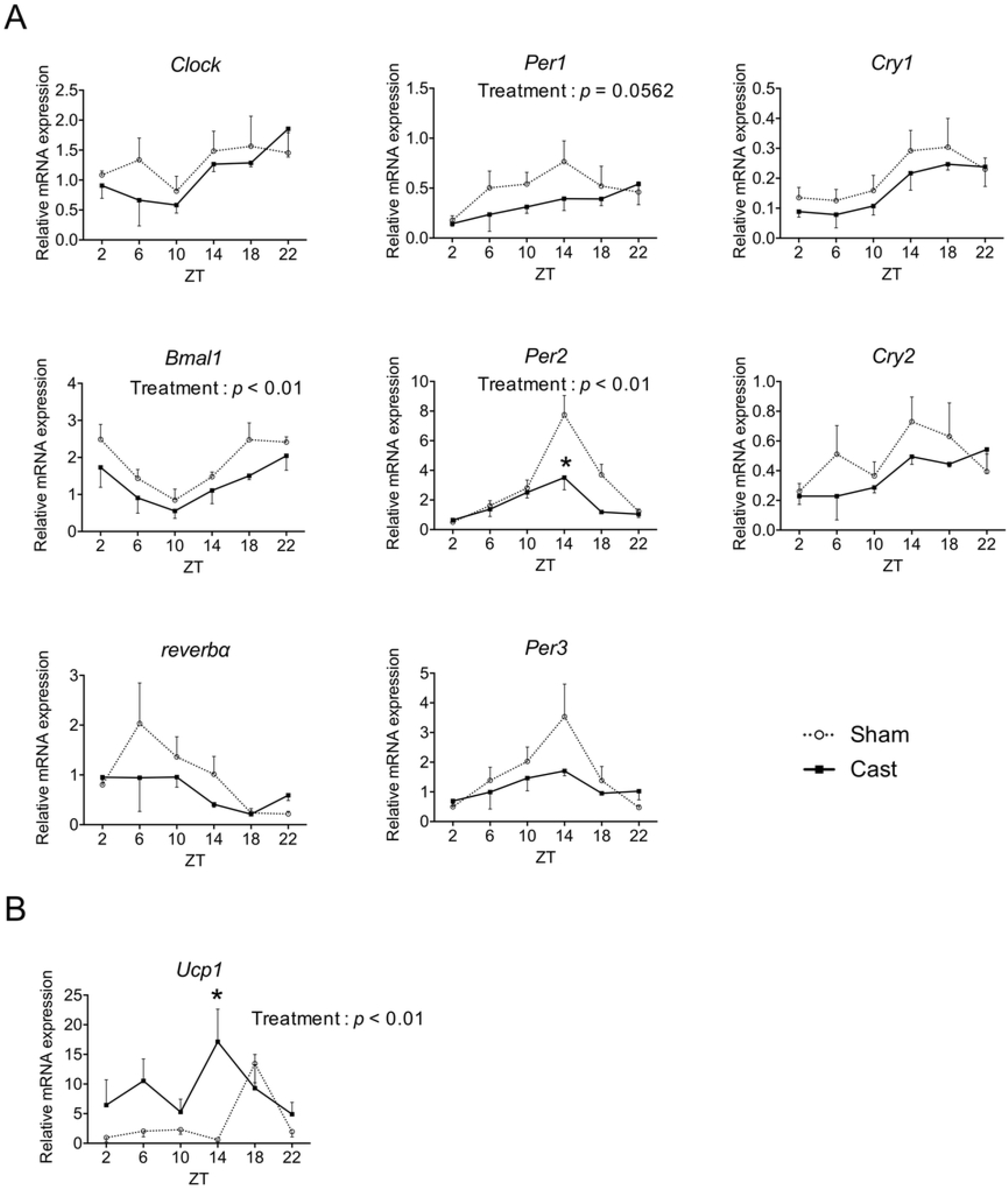
Circadian expression rhythms of clock genes and *Ucp1* in mouse scWAT. The relative expression of **(A)** clock genes (*Clock, Bmal1, Rev-erbα, Per1, Per2, Per3, Cry1, Cry2*) and **(B)** *Ucp1* was analyzed at ZT 2, 6, 10, 14, 18, and 22 by quantitative RT-PCR in scWAT of Sham (dot line) and Cast (solid line) mice (n = 4). The data are shown as the mean ± SEM. The main effect of treatment was detected by two-way ANOVA. * *p* <0.05 (Tukey’s multiple comparison test).

## Discussion

Metabolic disorders caused by castration are likely to occur in almost all animals, and studies have been conducted to elucidate the mechanism of their onset; however, these mechanisms have not been completely elucidated. We focused on thermogenic *Ucp1* as the cause of metabolic disorders and aimed to elucidate how castration causes abnormal expression of *Ucp1*. In this study, castrated mice were found to have a reduced rate of weight gain without any effects on food intake, locomotor activity, or muscle mass. It is well known that castrated mice show a reduction in weight gain that does not result from altered food intake on standard chow [7,24]. It has also been reported that although some castrated mice occasionally show increased or decreased behavioral activity, castration surgery on mice does not affect locomotor activity in the open field test [25– 28]. Moreover, castration of mice does not affect the weight of the gastrocnemius muscle [29]. Our castrated mice also demonstrated increased body temperature during the light phase, which is a resting period. Castration model animals are also known to have an improved ability to produce heat from BAT or scWAT and to suppress weight gain with increasing body temperature [23,30,31]. In support of this ability, our castration models had reduced epididymal fat weight and showed a fat combustion effect.

The mechanism of nonshivering thermogenesis resulting from the expression of *Ucp1* in BAT and scWAT is widely known in mice and humans [32,33]. In particular, the heat production function of the beige adipocytes in scWAT plays an important role in the prevention of obesity and metabolic diseases; however, it is not fully understood. A few previous studies have demonstrated that the expression of *Ucp1* in scWAT is increased in male rodents after gonadectomy [23,34]. As in previous reports, the expression levels of *Ucp1* significantly increased in the scWAT of castrated mice after cold stimulation at 4 °C for 4 hours compared with the expression in sham-operated mice, although there was no effect on the *Ucp1* expression in BAT. The reason for the increased *Ucp1* expression in scWAT without affecting expression in BAT is not known in detail, but it may be related to the shorter cold stimulation time compared to previous reports. Notably, the expression of the thermogenesis-related genes *Pparγ, Prdm16*, and *Pgc1-α* was not affected by castration in our study. Therefore, we hypothesized that castration induced *Ucp1* expression without being mediated by these thermogenesis-related genes. A recently published paper supports our hypothesis that activin E directly increases *Ucp1* expression in inguinal WAT without affecting the expression levels of these thermogenesis-related genes [35].

Previous reports have shown that castration causes a decrease in the androgen receptor in the SCN and that the biological rhythms fluctuate in these mice [18,36]. Recently, in another rodent model, the Syrian hamster, it was found that the expression of *clock* in the SCN is decreased after gonadectomy [37]. In this study, we also observed that castrated mice showed decreases in circadian behavioral rhythm and amplitude; moreover, the expression of *clock* in the SCN was also decreased. It has been suggested that testosterone deficiency caused by castration leads to disturbances in the rhythm control function of the SCN [18,36,38]. In our study, the disturbance of circadian rhythm due to castration may have been influenced by testosterone deficiency to some extent. On the other hand, castration is also known to affect peripheral clocks in addition to disturbing central rhythms. In castrated rodents, the expression of clock genes shows various circadian fluctuations in prostate tissue, limb skeletal muscle, and liver tissue compared with the expression in sham animals [19,39,40]. Similarly, in our animals, the expression rhythms of many clock genes in the peripheral scWAT also fluctuated with castration and were significantly reduced, especially those of *Bmal1* and *Per2*. Notably, the expression of *Ucp1* in scWAT also shows a circadian rhythm, but the expression level increases after castration, as opposed to the decreases in the levels of clock genes such as *Bmal1* and *Per2. Bmal1*-null (KO) mice exhibit a rhythm of high *Ucp1* expression, and *Ucp1* is highly expressed in the BAT of cold-exposed *Bmal1*-null mice, although different mechanisms are involved in the case of adrenaline stimulation [41]. Furthermore, *Bmal1* expression affects the TGFβ pathway and BMP signaling, and decreased *Bmal1* expression leads to improved heat production capacity due to high *Ucp1* expression in BAT [42]. Similar to the link between *Bmal1* and *Ucp1, Per2* is known to be deeply involved in the regulation of Ucp1 expression. Since *Per2* inhibits the binding of the *Ucp1* gene to the promoter region by phosphorylating *Ppar*γ2 s112, the expression of the *Ucp1* gene is suppressed under high expression of *Per2*, and *Per2* deficiency leads to high expression of the *Ucp1* gene specifically in WAT [43]. These past reports support our results. In addition to being regulated by *Bmal1* and *Per2* in BAT and WAT, several previous papers have shown that the clock gene is highly involved in the regulation of *Ucp1* expression. Using *Rev-erbα-*knockout mice, Gerhart-Hines *et al*. demonstrated that the nuclear receptor *Rev-erbα*, a clock gene, regulates the expression of *Ucp1* in BAT to establish and maintain a temperature rhythm suitable for environmental requirements [44]. It was also reported that in HIB1B preadipocyte cell cultures, the serotonin-regulated clock gene *Bmal1* plays an important role in the expression of *Ucp1* in brown adipocytes in the early stages of differentiation [45]. Moreover, although the detailed mechanism has not yet been fully elucidated, it is widely known that disturbance of the circadian clock leads to the development of metabolism-related disorders [46–48]. Together, these data suggest that castration not only disturbs the central circadian clock of mice but may also strongly affect the clock mechanism in peripheral scWAT, especially the rhythmic expression of *Bmal1* and *Per2*, which alters the expression of *Ucp1*.

## Conclusion

In conclusion, our data demonstrate that castration causes disturbances in the master clock, which disrupts clock function in scWAT, eliminates the order of clock gene-controlled *Ucp1* expression rhythm, and impairs thermogenic function. Further experiments will be needed on the detailed mechanism of how castration impacts clock gene expression. Our results will help to contribute to the improvement in metabolic diseases and quality of life (QOL) of castrated livestock, companion animals and humans.

## Acknowledgments

T.O. designed this study and wrote this paper. H.O. performed the experiment, and M.S. and M.M. assisted with the experiment. T.O. and H.O. analyzed the data. All members discussed the results and this manuscript.

